# Bacterial cell shape control by nutrient-dependent synthesis of cell division inhibitors

**DOI:** 10.1101/2021.03.25.436990

**Authors:** Nikola Ojkic, Shiladitya Banerjee

## Abstract

By analysing cell shape and size of the bacterium *Bacillus subtilis* under nutrient perturbations, protein depletion, and antibiotic treatments we find that cell geometry is extremely robust, reflected in a well-conserved scaling relation between surface area (*S*) and volume (*V*), *S* ~ *V^γ^*, with *γ* = 0.85. We develop a molecular model supported by single-cell simulations to predict that the surface-to-volume scaling exponent *γ* is regulated by nutrient-dependent production of metabolic enzymes that act as cell division inhibitors in bacteria. Using theory that is supported by experimental data, we predict the modes of cell shape transformations in different bacterial species and propose a mechanism of cell shape adaptation to different nutrient perturbations. For organisms with high surface-to-volume scaling exponent *γ*, such as *B. subtilis*, cell width is not sensitive to growth rate changes, whereas organisms with low *γ*, such as *A. baumannii*, cell shape adapts readily to growth rate changes.

**SIGNIFICANCE:** How bacteria regulate their size and shapes to optimise their growth fitness in different nutrient environments remains largely unknown. By analysing the surface area and volume of rod-shaped *B. subtilis* exposed to different nutrient conditions and antibiotics we find that cells preserve a power law scaling between surface area and volume. We show that the surface-to-volume scaling is extremely robust and is regulated by nutrient-dependent synthesis of cell division inhibitors. By analysing different bacterial types, we find that cells conserve the surface-to-volume scaling exponent that is typical for each species, implying distinct mechanisms for morphological adaptation in each organism.

---

Control of cell size and shape is essential for bacterial growth, proliferation, nutrient access, motility and adhesion (1). It is well known that bacterial morphologies are highly adaptive and varies with growth conditions (2-4). Studying the growth of *S. enterica* in different nutritional conditions, Schaechter *et al*. discovered the *nutrient growth law* (5) — cell size increases exponentially with growth rate. Recent single-cell studies have confirmed this result for evolutionary divergent *E. coli* and *B. subtilis* (6, 7). However, not all cell types change shapes the same way. While *E. coli* grows in both length and width (3, 6, 8), *B. subtilis* primarily elongates in length (7, 9). It is poorly understood how a bacterium may regulate cell length and width in response to growth conditions.

During steady-state growth, cell size is determined by a balanced trade-off between the rates of cell growth and division (10). One of the main molecular players that control bacterial division rate is the tubulin homolog FtsZ that triggers cell division when it reaches a certain threshold amount in the Z-ring (3, 11). FtsZ is produced in the cytoplasm and is subsequently recruited in the ring in a concentration dependent manner (3). How this assembly-competent pool of FtsZ is controlled in different nutrient conditions is key to understanding the cell size regulation in bacteria (10).

Weart, Levin and colleagues identified the metabolic sensor UgtP as the inhibitor of FtsZ ring assembly in *B. subtilis* (12). In this organism, nucleotide sugar (UDP-glucose) controls the affinity of UgtP to FtsZ, establishing a connection between nutrient conditions and division protein availability (13). In nutrient poor medium, UgtP mostly binds to itself without interfering with the FtsZ pool. However, in nutrient rich medium UgtP binding affinity to FtsZ drastically increases, leading to depletion of the assembly-competent pool of FtsZ. This results in delayed cell division and cell enlargement (13). Despite a change in UgtP affinity to FtsZ with sugar availability, it is unclear how the cytoplasmic pool of UgtP and its synthesis depend on the growth rate, which in turn regulates FtsZ assembly and cell size. While similar division inhibitors have been identified in other bacterial species (14–16), it remains unknown how these inhibitors are regulated to control cell morphology in different nutrient conditions.

To investigate cell shape control in *B. subtilis*, we first analysed population-averaged cell volume and surface area data in different nutrient conditions (Fig. 1A) (7). In contrast to *E. coli* cells where *S* ∝ *V*^2/3^, as a consequence of aspect ratio preservation (3), for *B. subtilis S* ∝ *V*^0.83±0.03^ and aspect ratio increases with growth rate (9) (Fig. 1A, C). This scaling relation is a consequence of marginal dependence of cell width on growth rate, while *S* ∝ *V* would imply that width stays constant (Fig. 1D). To interrogate the robustness of this scaling relation, we analysed surface area and volume data of cells treated with five different types of antibiotics that target membrane biosynthesis (AZ105, Cerulenin), cell-wall (D-Cycloserine, Mecillinam, Phosphomycin), RNA synthesis (rifampicin), and DNA (ciprofloxacin) (17). Under these perturbations, cells maintained the scaling relation *S* ∝ *V*^0.88±0,03^, identical to when cells are exposed to protein degradation systems mimicking targeted protein depletion by antibiotics. Further, when 289 essential proteins are depleted using CRISPRi (18), surface-to-volume scaling stayed surprisingly constant and extremely robust (*S* ∝ *V*^0.855±0.005^). Even though there is a correlation between median cell length and width for cells growing at steady-state (18), median cell width changes marginally from 0.92 to 1.16 *μ*m in comparison to larger changes in median cell length between 3.5 and 12.7 *μ*m. This coupling between cell width and length follows from the scaling relation *S* ∝ *V*^0.85^. Normalising all measurements with the smallest corresponding values of surface area and volume, all data collapsed to a single curve demonstrating the universality of 0.85 scaling in *B. subtilis* under nutrient, antibiotic or genetic perturbations (Fig. 1A, inset).

**Figure 1:**
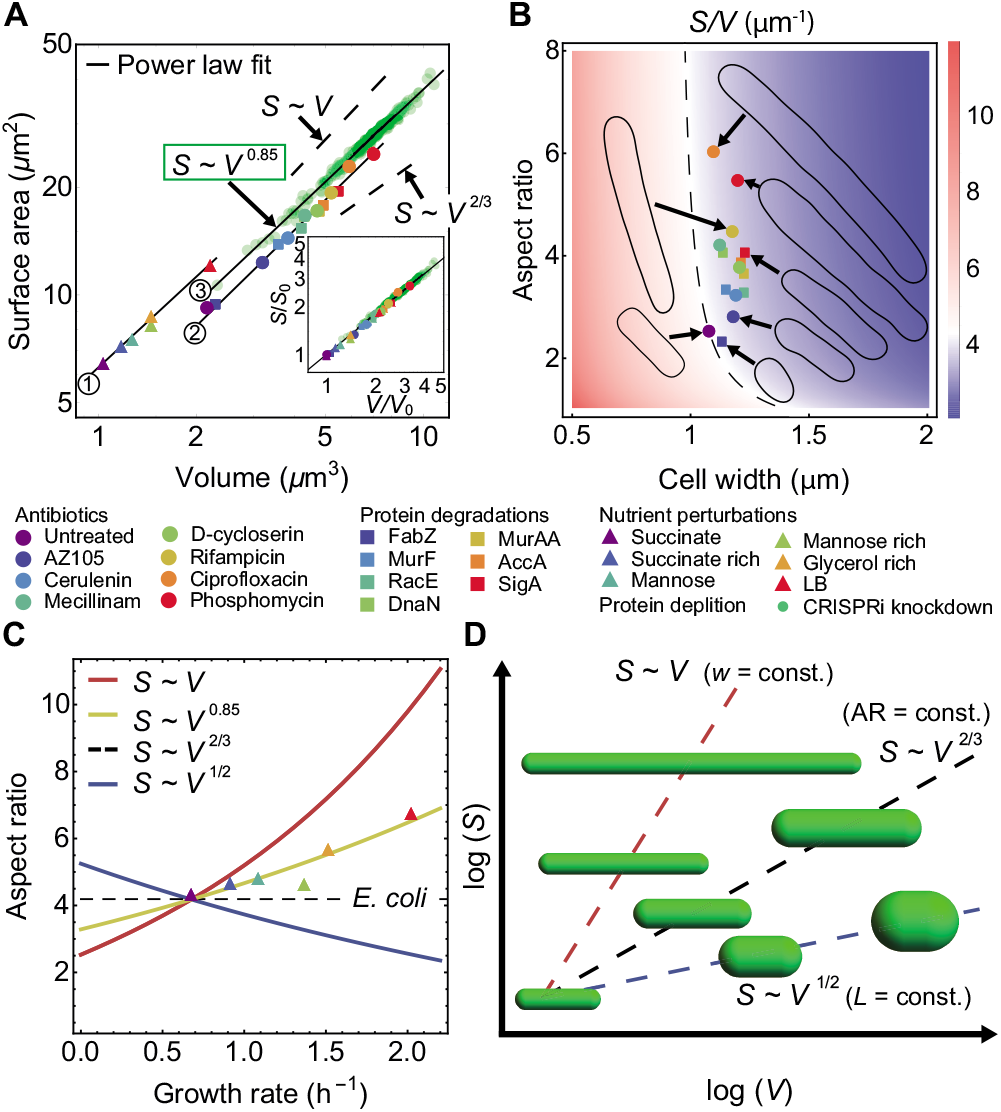
Surface-to-volume scaling and cell shape control in *B. subtilis*. (A) *B. subtilis* cells exposed to different antibiotics, nutrient conditions, protein degradation and depletion (7, 17, 18) conserve the scaling relation between population averaged surface area (*S*) and volume (*V*): *S* ∝ *V*^0.85^. Scaling exponent for three different background strains shown with circled number are 0.88 ± 0.03, 0.83 ± 0.03, and 0.855 ± 0.0.05. (Inset) Surface area vs volume data, normalized by the corresponding smallest quantities, collapse on the curve *S* ∝ *V*^0.85^. (B) Surface-to-volume ratio (*S/V*) decreases for cells exposed to different antibiotics or protein degradation. Untreated cells have *S/V* ≈ 4.2 *μ*m^-1^. Typical cell contours under perturbations are shown as inset. Dashed black line is constant *S/V* for unperturbed cells. (C) Aspect-ratio (length divided by width) vs growth rate for cells grown in different nutrient conditions (points); theoretical predictions from growth law (Fig. S1A) for different surface-to-volume scaling exponents (lines), without further fitting. (D) *S-V* scaling for different cell shapes showing the modes of cell shape changes.

It was previously reported that Gram negative *E. coli, C. crescentus* and Gram positive *Listeria* decrease *S/V* with increasing nutrients and applied antibiotic stress (2). We therefore inquired how different antibiotics or genetic perturbations affect *S/V* in *B. subtilis.* Interestingly, for all analysed conditions *B. subtilis* adopts a lower surface-to-volume ratio compared to the untreated cell (Fig. 1B). This results from an increase in cell width and volume under stress. To understand the dependence of cell shape on the growth rate of the medium, we combined the nutrient growth law (*V* = *V*_0_ e^*αk*^) (Fig. S1A) and the scaling relation *S* = *μV^γ^* to derive a relation between cell aspect ratio (*a*) and growth rate, where *α, μ* and *V*_0_ are constants, *k* is the growth rate, and *γ* is the scaling exponent. Assuming a spherocylindrical geometry, appropriate for rod-shaped bacteria, we get:

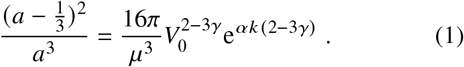

Solution to Eq. (1) predicts a non-linear relation between cell aspect ratio and growth rate for different scaling exponents (Fig. 1C). Consistent with our previous findings (3), the above relation predicts a constant aspect ratio of ≈ 4 for *γ* = 2/3 as observed for *E. coli.* Taken together, *B. subtilis* maintains a fixed scaling exponent *γ* = 0.85 under various growth perturbations, with cellular aspect ratio increasing with nutrient availability. Maintenance of a constant *γ* across 310 different conditions implies a highly conserved mode of shape changes in *B. subtilis*, reflecting intrinsic robustness of the molecular pathways that regulate cell shape and size.

To explain how nutrient availability regulates cell size and aspect ratio, we implemented a molecular model for cell division and size control in *B. subtilis*. In this model, a *B. subtilis* cell grows exponentially in length *L* at a rate *k* (d*L*/d*t* = *kL*), accumulates division proteins FtsZ in the cytoplasm, assembles a Z-ring in the mid-cell from the cytoplasmic FtsZ pool, and divides when a threshold abundance of FtsZ accumulates in the ring (Fig. 2A-D). Cytoplasmic FtsZ interacts with the metabolic enzyme UgtP that prevents Z-ring assembly in a nutrient-dependent manner, thereby delaying division. Dynamics of FtsZ are described using a three-component model (Fig. 2A). First, cytoplasmic FtsZ molecules with abundance *F*_c_ grows at a rate proportional to cell volume, such that the amount of accumulated FtsZ is proportional to the added cell size in one division cycle (3, 6, 10). Second, a ring-bound FtsZ component with abundance *F*_r_ is assembled from the cytoplasmic monomeric pool at a constant rate *k*_on_, and disassembled from the ring with a rate *k*_off_. Third, a component of the cytoplasmic FtsZ is sequestered by UgtP, resulting in the formation of UgtP-FtsZ heterodimer that does not associate with the ring (13). Dynamics of the system are described by the following coupled differential equations:

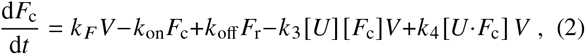

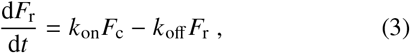

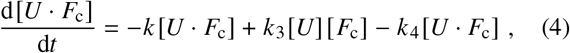

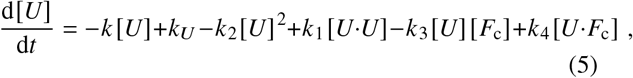

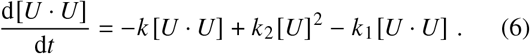

**Figure 2:**
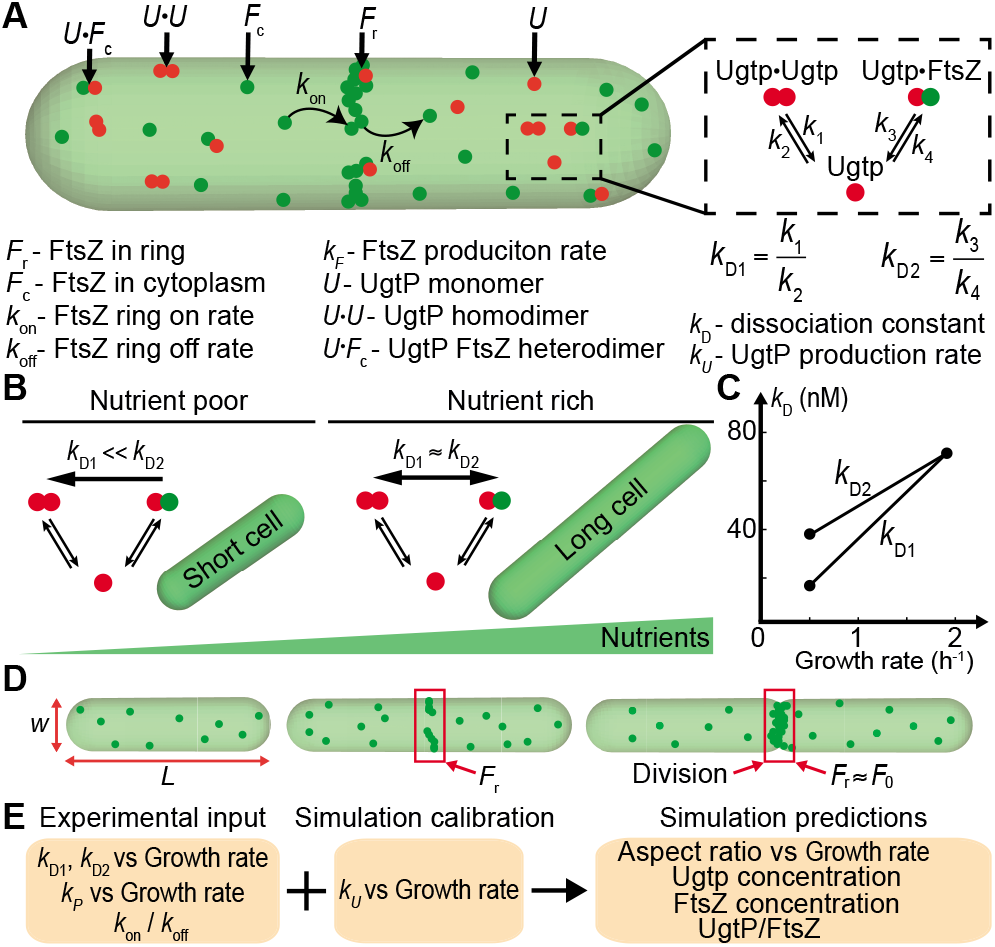
Molecular model for cell division and size control in *B. subtilis*. (A) Schematic of FtsZ dynamics in *B. subtilis.* FtsZ in the ring (*F_r_*) is recruited from a cytoplasmic pool of freely available FtsZ monomers (*F*_c_) with association rate *k*_on_ and is disassembled at a rate *k*_off_. Dashed box on the right: Cytoplasmic FtsZ is sequestered by UgtP creating *U* · *F*_c_ heterodymer with dissociation constant *k*_D1_ = *k*_1_/*k*_2_. UgtP also forms homodymer *U* · *U* with dissociation constant *k*_D2_ = *k*_3_/*k*_4_ (13). (B) Depending on the nutrient quality, UgtP has different binding affinity to itself and to FtsZ. In nutrient-poor conditions, most UgtP binds to itself forming homodimers, while in nutrient-rich environment UgtP sequesters FtsZ, preventing FtsZ recruitment in the ring and delayed cell division. (C) Dependence of *k*_D1_ and *k*_D2_ on growth rate (13). (D) Cell cycle of single bacterial cell of length (*L*), and width (*w*). When FtsZ amount in the ring (*F*_r_) reaches a threshold *F*_0_, cell division is triggered. (E) Experimental inputs for model parameters are integrated in simulations. We calibrate *k_U_* to reproduce experimental data for aspect ratio vs growth rate.

In the above, *k_F_* is the production rate of cytoplasmic FtsZ, *k_U_* is the UgtP production rate, *k* is the cell growth rate, *k*_D1_ = *k*_1_/*k*_2_ is the UgtP homodimer dissociation constant, and *k*_D2_ = *k*_4_/*k*_3_ is UgtP-FtsZ heterodimer dissociation constant. Assuming balanced biosynthesis (3, 6), *k_F_* is taken to be proportional to *k*. In Eqs. (4)-(6), the first terms on the right hand side represent protein dilution due to cell growth.

In the above model, the availability of free and sequestered FtsZ changes across nutrient conditions due to growth rate dependence of the dissociation constants, *k*_D1_ and *k*_D2_. Based on experimental findings (13), both *k*_D1_ and *k*_D2_ increase with nutrient availability (Fig. 2B). The increase in dissociation constants with growth rate shifts the chemical equilibrium from mostly free FtsZ available for ring recruitment in nutrient poor medium, to partially sequestered FtsZ that are unable to build the ring in nutrient rich medium (Fig. 2B). Since experimental measurements of dissociation constants were only available for slow and fast growth conditions (13), we assumed a linear interpolation of the growth-rate dependence of the dissociation constants between the two measured values (Fig. 2C). Our simulation results are robust to the assumption of a linear interpolation function for *k_D_* (Fig. S2). Since *k_F_* is taken proportional to growth rate *k* across nutrient conditions, FtsZ concentration stays unchanged (Fig S2B), as experimentally observed (13). In addition, the ratio of the *on* and *off* rates for ring assembly is taken to be constant, *k*_on_/*k*_off_ = 1/2, so that ≈ 30% of FtsZ is recruited in the ring, independent of growth rate (19). Similar fraction of FtsZ recruitment in the ring was also reported in *E. coli* (11, 20).

In our simulations, cell division occurs when the amount of FtsZ recruited in the ring (*F*_r_) reaches a threshold *F*_0_ (3,11) (Fig. 2D), to satisfy the adder principle for cell size homeostasis (21). It was previously proposed by us (3) and later experimentally shown (22) that the threshold amount depends on cell width, *F*_0_ ∝ *w*. Control of cell width follows from the model (2): d*S*/d*t* = *βV*, where *β* is the surface synthesis rate that is linearly proportional to growth rate in *B. subtilis* (Fig S1B). As a result, cell width is almost unperturbed across nutrient conditions and *F*_0_ stays approximately constant.

We numerically simulated Eq. (2)-(6) in 10^5^ asynchronous cells to determine the dependence of population-averaged cell length and aspect ratio on growth rate. In simulations we always assumed a binary fission even for the long cells (23, 24). For each value of growth rate, we calibrated the UgtP production rate *k_U_* to fit the experimental data for aspect ratio vs growth rate in the case *S* ∝ *V*^0.85^ (Fig. A, B). The best fit *k_U_* increases non-linearly with growth rate (Fig. 3B, third curve from bottom), such that UgtP monomer amount increases from ≈ 1100 in slow media to ≈4600 in fast growth conditions (Fig. S3A), similar in magnitude to estimates made from experimental data (13). A nonlinear increase in *k_U_* with *k* implies an increased rate of FtsZ inhibition by UgtP (∝ [*U*] ≈ *k_U_*/*k*) in fast growth conditions, leading to longer cells. By calibrating the dependence of *k_U_* on *k* for different shape families (*S* ∝ *V^γ^*), we predict the variation in aspect ratio with growth rate (Fig. 3A). A linear increase in *k_U_* with *k* predicts a constant aspect ratio (*S* ∝ *V*^2/3^), whereas a decrease of *k_U_* with *k* leads to aspect ratio decreasing with growth rate (*S* ∝ *V*^1/2^).

**Figure 3:**
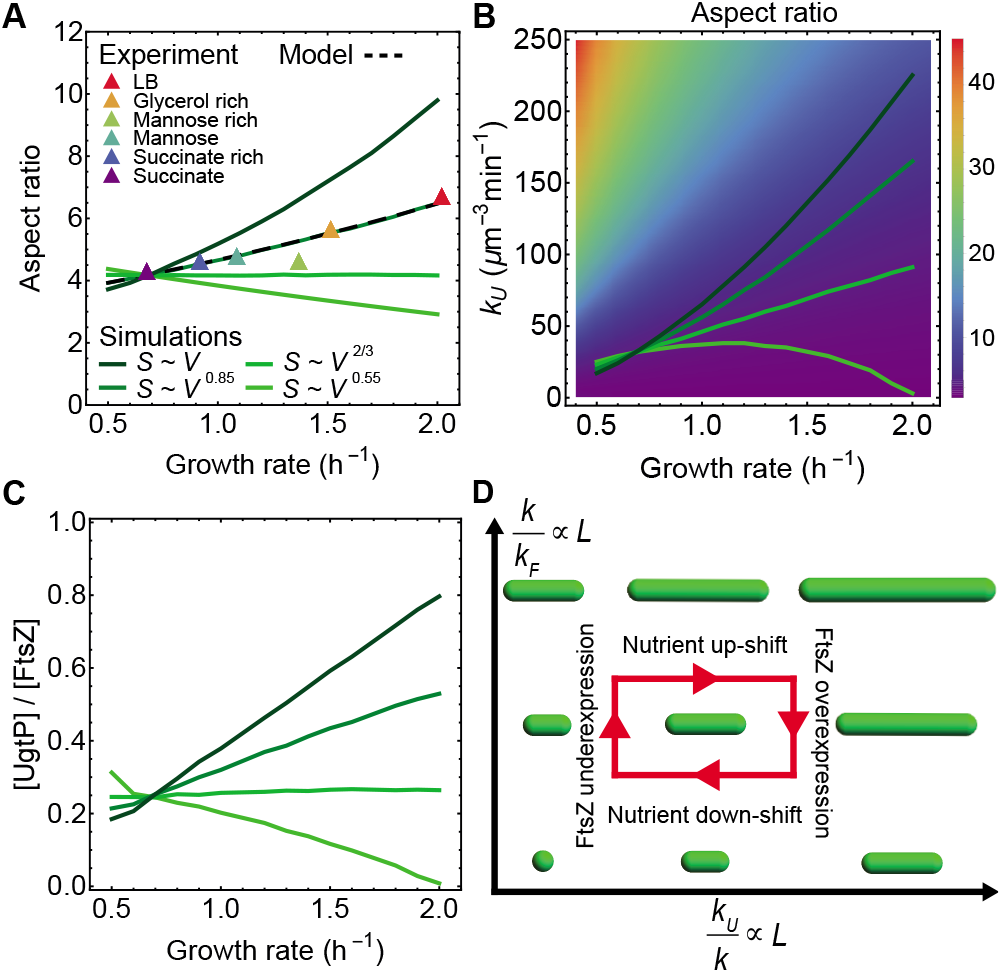
Model predictions for *B. subtilis* cell shape and size in different nutrient conditions. (A) Simulation results for the dependence of aspect ratio on growth rate in four different scaling regimes. UgtP production rate (*k_U_*) was calibrated parameter to fit experimental data for the dependence of aspect ratio on growth rate (*k*) (7). (B) Population-averaged aspect ratio of cells for different values of *k* and *k_U_*. The solid lines represent the predicted dependence of *k_U_* on *k* to reproduce the aspect-ratio dependence on growth rate in panel A. (C) Simulation predictions for the ratio of the concentration of UgtP and FtsZ as a function of growth rate. (D) Phase diagram for cell shape changes as function of *k*/*k_F_* and *k_U_/k*.

Our model successfully captures that the ratio of UgtP to FtsZ concentration increases with nutrient quality, with [UgtP]/[FtsZ]≈0.5 in fast growing conditions (Fig. 3B), in accord with experimental estimates (13). Even though FtsZ monomer amount increases with nutrients, FtsZ concentration stays approximately constant ≈ 9 *μ*M (Fig. S3B), as observed experimentally and consistent with reported estimates of FtsZ concentration range 4-10 *μ*M (12, 13). Despite an increase in the total number of FtsZ monomers with nutrients, the fraction of ring assembly-competent monomers decreases, leading to slower ring assembly and longer cells (Fig. S3C).

By systematically varying the rate of growth (*k*), FtsZ synthesis (*k_F_*) and UgtP synthesis (*k_U_*) across nutrient conditions (Fig. 3D, S4), we predict two fundamental modes of cell morphological changes. First, by increasing *k_U_*/*k*, the abundance of ring assembly-competent monomers decreases, giving rise to elongated cell morphologies. Second, by increasing *k/k_F_*, cells increase in length as the rate of production of division proteins is slower than the rate of cell elongation (10).

Since division inhibitors are not unique to *B. subtilis* and have been identified in other bacterial species (14-16), we ask how the synthesis of division inhibitors can be regulated to control the cell morphology. One of the key predictions of our simulations is that there is a one-to-one correspondence between the scaling exponent *γ* and how fast the inhibitors are produced relative to growth rate. If *k_U_* increases linearly with *k*, then *γ* = 2/3 and the aspect ratio remains constant, whereas if *k_U_* decreases with *k* then *γ* < 2/3 and aspect ratio decreases with *k*. In general, *γ* increases monotonically with d*k_U_*/d*k* (normalised by its value at *γ* = 2/3) (Fig. 4A). Since different organisms maintain different values of *γ* (Fig. 4B,S4), we predict that they have distinct regulatory functions for modulating *k_U_* in response to changing nutrient conditions.

**Figure 4:**
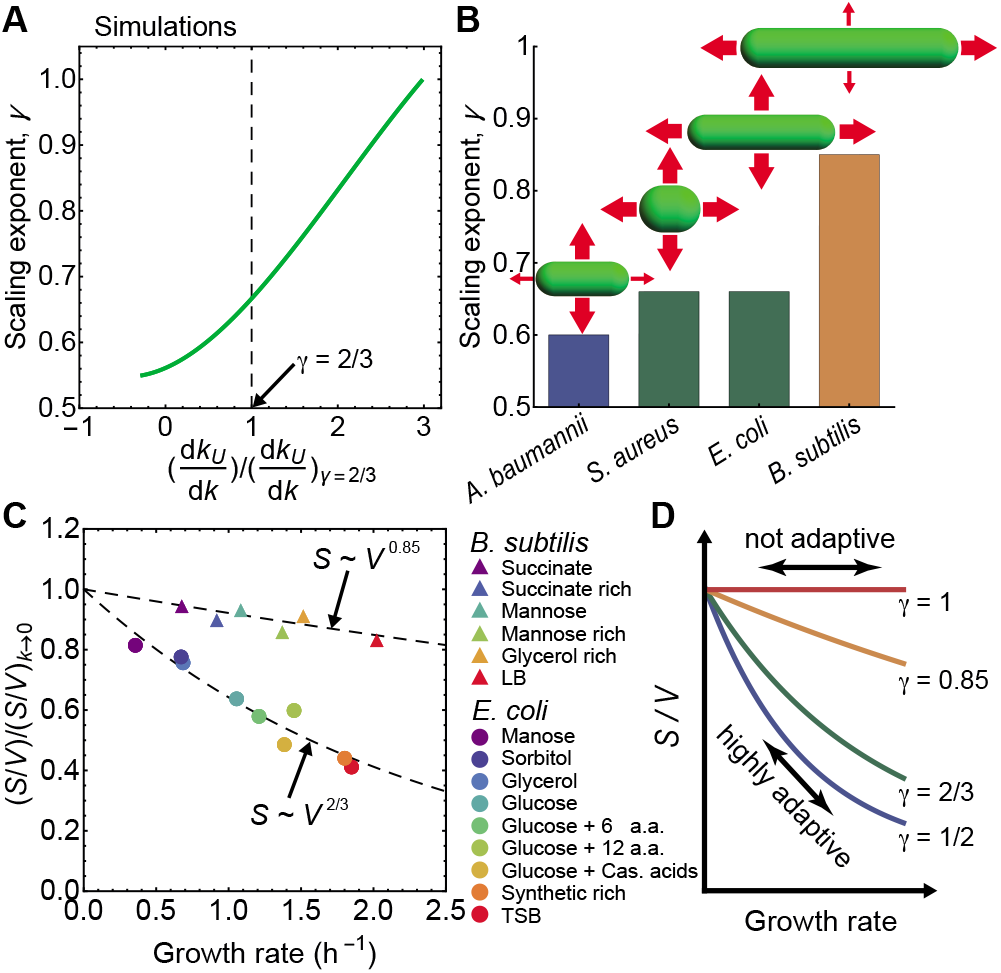
Surface-to-volume scaling and cell shape adaptation in different bacterial species. (A) Model prediction for surface-to-volume scaling exponent *γ* as a function of d*k_U_*/d*k*: rate of increase in UgtP synthesis per unit increase in growth rate. (B) Scaling exponent for different organisms (see Fig. S5): *A. baumannii* (25), *S. aureus* (26), *E. coli* (see (3)) and *B. subtilis* (see Fig. 1A). Red arrows indicate the modes of cell shape changes. (C) Dependence of *S/V* on growth rate for *E. coli* and *B. subtilis.* Dashed lines are theoretical fits. (D) Variation of *S/V* with growth rate for different values of *γ*. If *S* ∝ *V* then cell shape is invariant to nutrient changes. If instead *S* ∝ *V*^1/2^ cells readily adapt *S/V* to growth rate.

Surface-to-volume scaling exponent constrains the modes of cell shape changes (Fig. 4B). Combining *S* = *V^γ^* with the nutrient growth law predicts how *γ* regulates cell shape changes in response to growth conditions. While *S/V* remains invariant for *γ* = 1, it increases with decreasing growth rate for *γ* < 1, such that a lower *γ* leads to larger *S/V* changes (Fig. 4C-D). Increase in *S/V* with lowering growth rate has an adaptive benefit that increases nutrient access through the cell surface in nutrient-poor medium. This may provide clues to why bacteria are observed to maintain *γ* < 1 across species.

## AUTHOR CONTRIBUTIONS

N.O. and S.B. designed the research. N.O. carried out simulations and analyzed the data. N.O. and S.B. wrote the article.

## ACKNOWLEDGMENTS

We gratefully acknowledge funding from EPSRC (EP/R029822/1), and Royal Society (URF/R1/180187, RGF/EA/181044).

