## Supplemental Material for "Bacterial cell shape control by nutrient-dependent synthesis of cell division inhibitors"

**N. Ojkic and S. Banerjee**

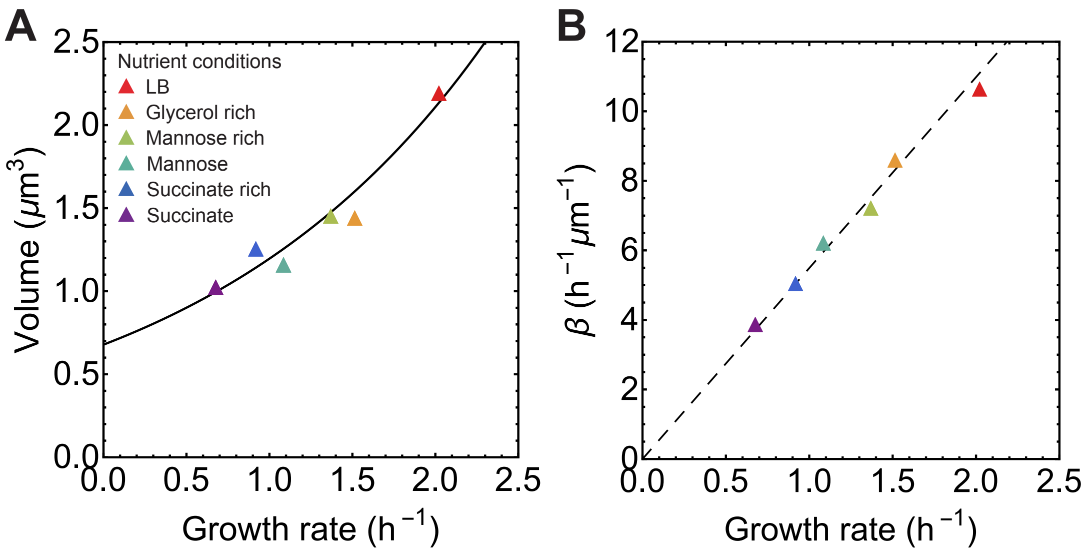


**Figure S** 1. **Growth-rate dependence of cell volume and surface area synthesis in *B. subtilis***. (A) Nutrient growth law — bacterial cell volume ($V$) vs growth rate ($k$). Best fit shown with black line: $V = (0.68 \pm 0.07)\mathrm{e}^{(0.56\pm0.07)k}$. (B) Surface area synthesis rate $\beta$ vs growth rate $k$, with $\beta = 4k/w$ [1, 2]. Here $w$ is the average cell width for a given nutrient condition. Best fit line is shown by the dashed black line: $\beta = (5.50 \pm 0.10)k$. All data obtained from [3].

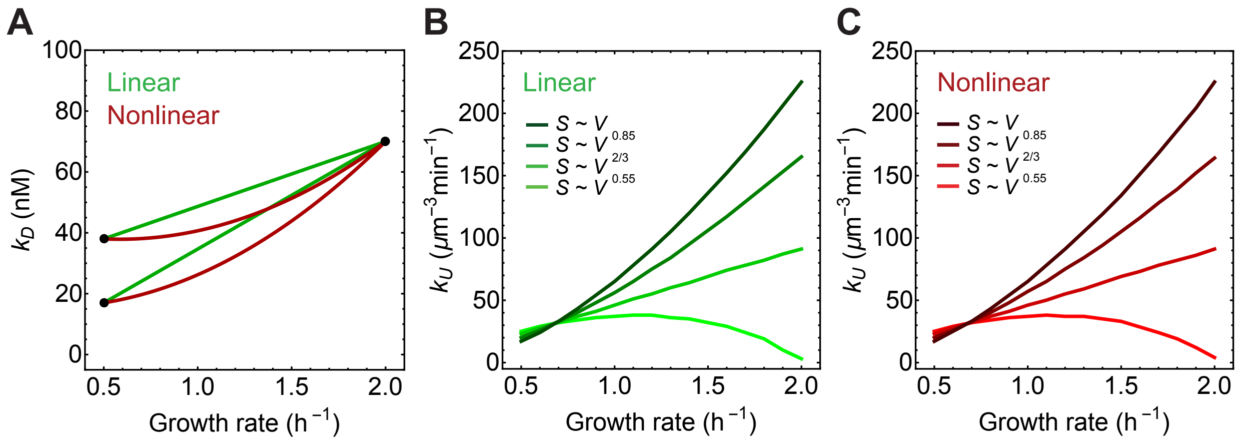


**Figure S 2. Robustness of simulation results under linear or nonlinear variation of $k_D$ with $k$.** (A) Linear (green) and nonlinear (red) interpolation of the dependence of $k_D$ on $k$. (B) $k_U$ vs $k$ for linear assumption (same as Fig. 3B). (C) $k_U$ vs $k$ for nonlinear interpolation, showing that the interpolating function does not affect the predicted dependence of $k_U$ on $k$.

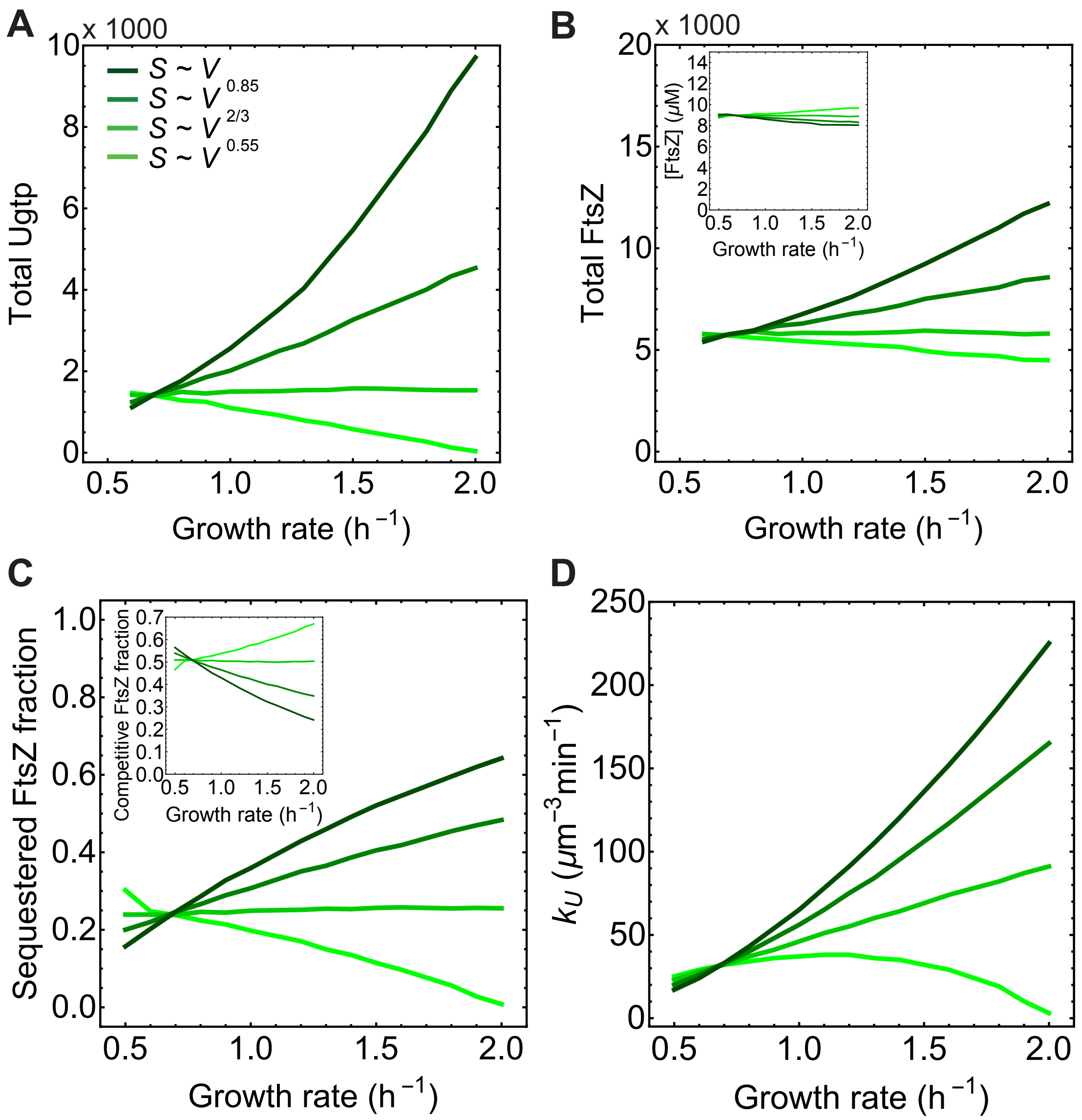


**Figure S** 3. **Single-cell simulations predict dynamics of division protein FtsZ and division inhibitor UgtP**. (A-B) Simulation predictions for total amounts of UgtP and FtsZ as function of growth rate for four different scaling relations between surface area and volume. (C) Fraction of sequestered FtsZ ($U \cdot F_c$) and total FtsZ vs growth rate. (Inset) Fraction of ring assembly-competent FtsZ ($F_c$) and total FtsZ vs growth rate. (D) UgtP production rate ($k_U$) vs growth rate that was calibrated to reproduce experimental data for aspect ratio vs growth rate ($k$) shown in Figure 2A.

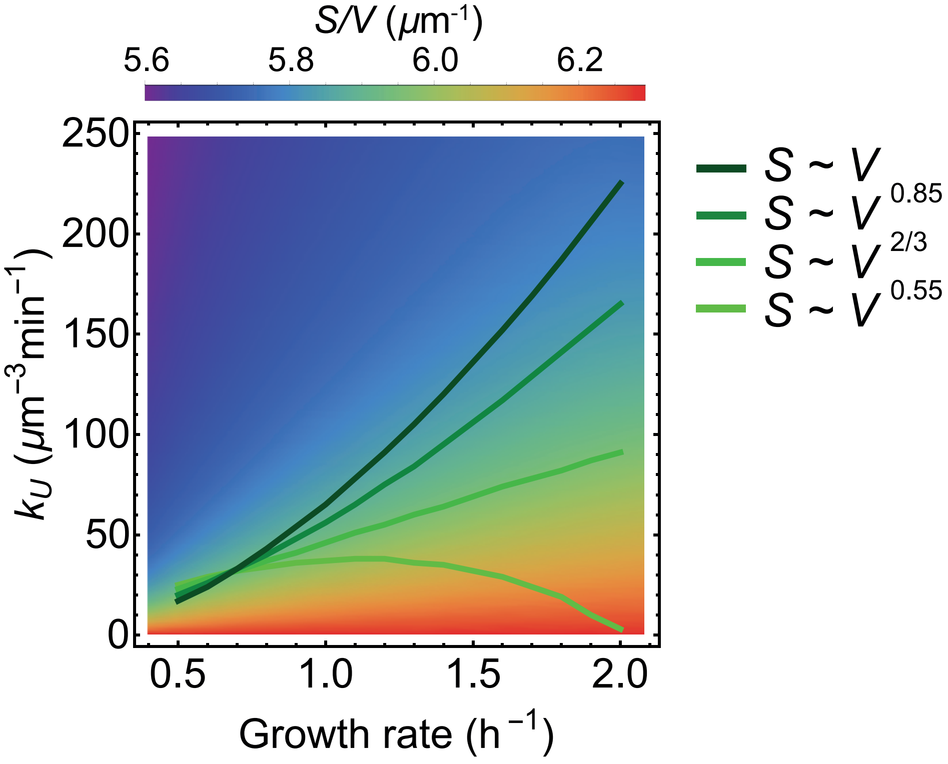


**Figure S 4. Simulation predictions for cell surface-to-volume ratio.** Population average of surface-to-volume ratio ($S/V$) of asynchronous steady-state cells for different values of growth rate ($k$) and UgtP production rates ($k_U$). The four curves for $k_U$ vs $k$ correspond to the four different scaling models for cell shape shown on the right.

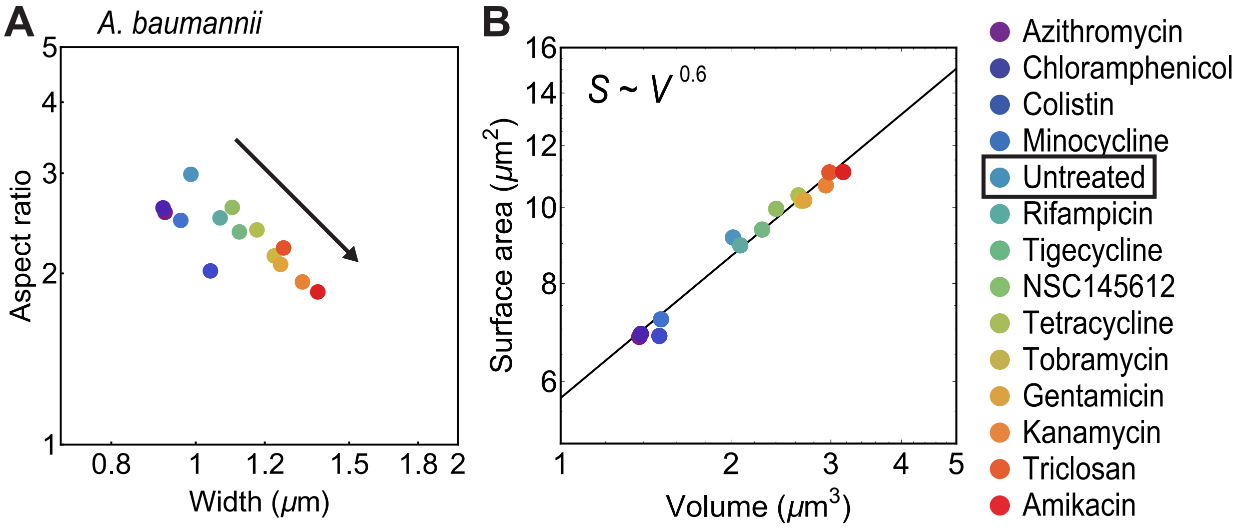


**Figure S 5. Morphological changes in *A. baumannii* under antibiotic treatment**. (A) Population averaged aspect ratio vs cell width for different antibiotic treatments shown on the right. Aspect ratio decreases with increasing width shown with black arrow. (B) Cell surface area ($S$) vs volume ($V$) for different antibiotic exposure. Black line is best fit: $S = (5.72 \pm 0.14)V^{0.60\pm0.02}$. All data obtained from [4].
